# A coordinated haptic mechanism ensures efficient DNA sampling by the 8-oxoguanine glycosylase OGG1

**DOI:** 10.1101/2025.10.08.681224

**Authors:** Maximilien Martinez, Ostiane D’Augustin, Despoina-Maria Chousianiti, Virginie Gaudon, Ana Gil Quevedo, Nicolas Bigot, Catherine Chapuis, Anne-Marie Di Guilmi, Jean-François Hubert, Bertrand Castaing, Anna Campalans, Sébastien Huet

**Affiliations:** Univ Rennes, CNRS, IGDR (Institut de Génétique et Développement de Rennes) - UMR 6290, BIOSIT (Biologie, Santé, Innovation Technologique de Rennes) - UMS 3480, US 018, F-35000 Rennes, France; Université de Paris-Cité, CEA/IBFJ/IRCM. UMR Stabilité Génétique Cellules Souches et Radiations, F-92260 Fontenay-aux-Roses, France; Université Paris-Saclay, CEA/IBFJ/IRCM. UMR Stabilité Génétique Cellules Souches et Radiations, F-92260 Fontenay-aux-Roses, France; Centre de Biophysique Moléculaire, UPR4301 CNRS, 45071 Orléans cedex 2, France; Graduate School « Santé, Science Biologique & Chimie du Vivant » (ED549), Université d’Orléans, France

## Abstract

7,8-Dihydro-8-oxoguanine (8-oxoG) is the most frequent base modification occurring upon oxidative stress. This highly mutagenic lesion is specifically recognized and excised by the DNA glycosylase OGG1 when paired with cytosine, initiating the base excision repair pathway. Since 8-oxoG neither significantly impacts the structure of the double helix nor blocks transcription and replication processes, its detection requires a careful inspection of each base pair by OGG1. By monitoring this lesion search process both *in vitro* and in living cells, we demonstrate that it involves a tight coordination between several conserved amino acids encircling the DNA helix. More specifically, we show that the N149–151 motif, on the target strand, as well as residues R154 and R204, on the opposite strand, both regulate OGG1 engagement on the DNA to ensure fast Y203-mediated base unstacking, a prerequisite for efficient 8-oxoG detection. These findings highlight the early mechanisms that enable OGG1 to maintain rapid sampling kinetics while preserving high specificity for 8-oxoG in the context of the complex architecture displayed by the DNA within the cell nucleus.

## INTRODUCTION

Our DNA is constantly exposed to reactive oxygen species (ROS) originating from cellular metabolism or exogenous sources. Among the wide range of DNA lesions generated by ROS, guanine oxidation is particularly prominent due to its low redox potential, leading to the formation of 7,8-dihydro-8-oxoguanine (8-oxoG)^1^. This oxidized form of guanine displays a strong mutagenic potential due to its ability to form a Hoogsteen base pair with an adenine in a *syn-anti* conformation, thus promoting G:C to T:A transversion^2,3^. Accumulation of 8-oxoG in the genome has been associated with various cancers and other age-related diseases such as neurodegenerative disorders^4^. The base excision repair (BER) pathway is the main route for eliminating modified bases in organisms ranging from bacteria to humans^5^. In mammals, 8-oxoG paired with a cytosine is specifically recognized and excised by 8-oxoguanine DNA glycosylase 1 (OGG1), thus initiating the BER pathway^6^. Besides this early role in the BER, OGG1 is also thought to contribute to the inflammatory response of the cell. Indeed, upon oxidative stress, OGG1 was shown to recruit to 8-oxoG containing inflammatory gene promoters to act as a scaffold promoting the accumulation of transcription factors such as NF-κB^7,8^.

Like any other factor in charge of detecting specific DNA alterations, OGG1 faces a difficult challenge due to the compact multiscale architecture of chromatin within the cell nucleus. Yet, in contrast to many other lesions, 8-oxoG neither induces double helix distortion nor blocks the transcription and replication machineries, two cues often used to detect the presence of a DNA lesion^9,10^. Therefore, OGG1 needs to carefully examine each base pair along the double-helix to efficiently detect and excise 8-oxoG. Single-molecule data on naked DNA demonstrated that OGG1 searches for its cognate lesion by transient scanning along the DNA^11–13^. During this scanning process, the intrahelical insertion of the wedge residue Y203 was proposed to promote base unstacking, a step that initiates the extrusion of the base from the double-helix for further inspection and potential cleavage^14–16^. Nevertheless, recent structural data did not reveal such an unstacked state for the OGG1/G:C complex and showed that Y203 does not directly contact the bases and rather interacts with N149^17^. These findings implied a kinetic discrimination between G and 8-oxoG, the latter being stabilized in an extrahelical conformation in contrast to the former. Yet, it remains unclear how Y203 and neighboring conserved OGG1 residues tune the kinetics of the elementary steps of this discrimination process, thus determining its specificity. Furthermore, several reports also demonstrated that OGG1 search mechanism is impacted by the wrapping of the DNA along histone octamers, questioning how molecular mechanisms observed along naked DNA translate in the context of the complex conformation displayed by chromatin^18–20^.

In the current work, we set up a multiscale analysis pipeline to study OGG1 behavior during the lesion-seeking process both *in vitro* and in living cells. Using this integrated framework, we demonstrate how a coordinated action of several conserved residues, Asn 149, 150 and 151 on one hand and Arg 154 and 204 on the other hand, ensures an efficient sampling of the DNA needed for the rapid and specific clearing of 8-oxoG within the nucleus.

## RESULTS

### OGG1 exploration of the nucleus involves transient binding to DNA associated with base unstacking

In order to better characterize the early stages of 8-oxoG clearance, we monitored the behavior of EGFP-tagged OGG1 in HeLa cells knocked-out (KO) for endogenous OGG1 using live cell fluorescence imaging. In line with previous findings, we observed that OGG1 is rapidly recruited after irradiation with a pulsed laser at 800 nm (Figure 1A), which was previously shown to induce 8-oxoG lesions^21^. This accumulation peaks at ∼10s post damage and is followed by a slower release of OGG1 that was previously showed to depend on the progressive excision of 8-oxoG^21,22^. This fast recruitment implies that OGG1 is able to efficiently find lesions buried within the DNA double helix. To get further insights regarding the OGG1 search mechanism, the motions of OGG1-EGFP within the nucleus in the absence of exogenous lesions was analyzed using fluorescence correlation spectroscopy (FCS). By estimating the residence time spent by the fluorescently tagged protein within the focal volume, we found that OGG1-EGFP was diffusing ∼10 times slower than a EGFP tandem whose molecular mass is similar to the tagged glycosylase (Figure 1B,C). These data agree with our previous observations^21^ and imply that OGG1 transiently associates with the DNA for base sampling while navigating within the nucleus. Given that 8-oxoG lesions do not distort the double helix structure nor block transcription and replication processes, this sampling relies on a careful inspection of the bases by OGG1. We analyzed this inspection process *in vitro* by monitoring the association between OGG1 and undamaged oligos using stopped-flow fluorescence spectroscopy. Upon mixing a double-stranded oligo bearing the fluorescent nucleobase analogue 2-aminopurine (2-aPu) facing a cytosine with purified OGG1, we observed a rapid fluorescence increase corresponding to the unstacking of the 2-aPu probe from the double-helix (Figure 1D,E), in line with previous observations^23^. These results show that OGG1 binding to DNA distorts of the local structure of the double-helix independently of the presence of its cognate lesion, possibly accompanied by a partial base extrusion as previously suggested^17^.

**Figure 1.**
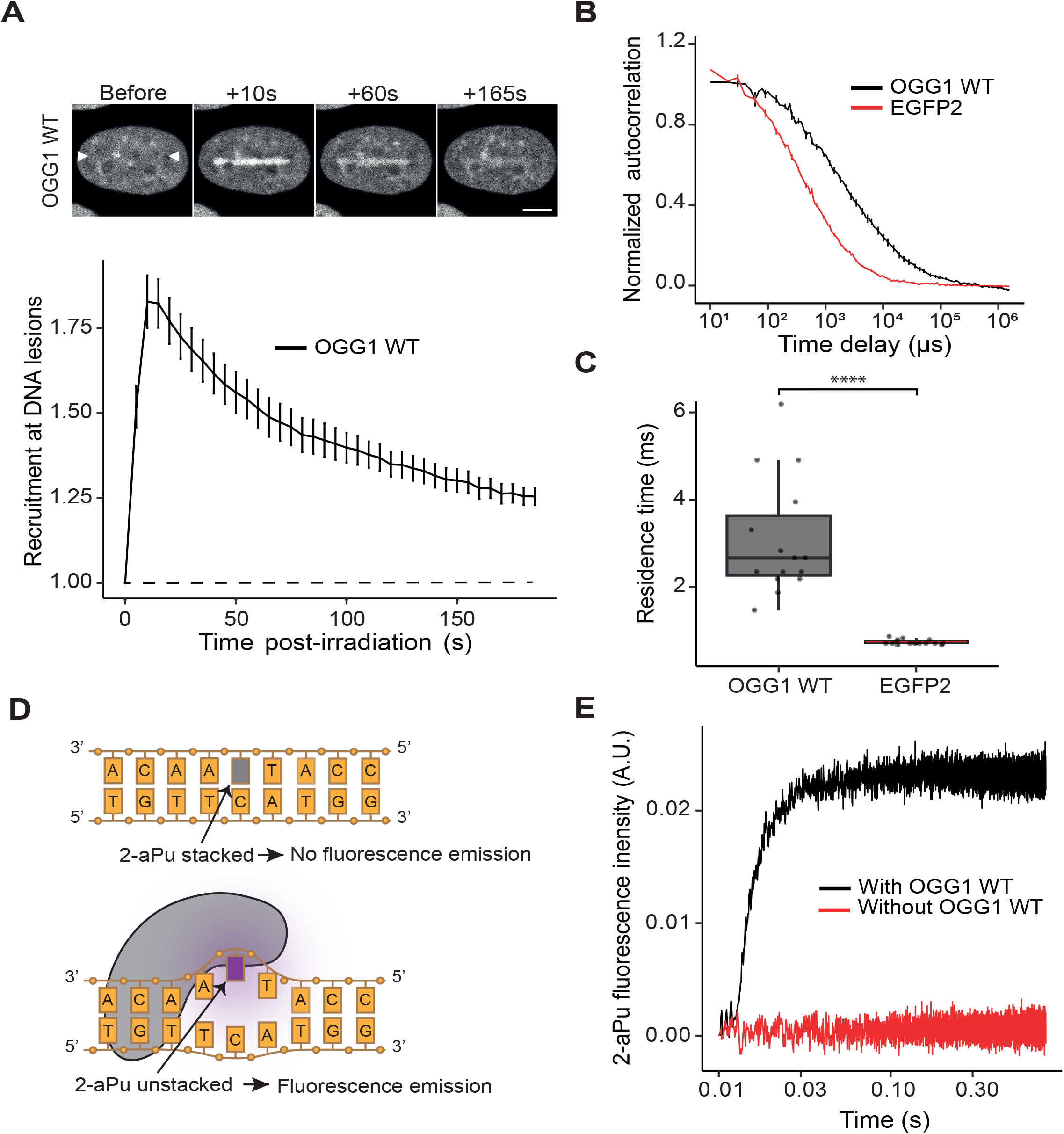
OGG1 exploration of the nucleus involves transient binding to DNA associated with base unstacking. **(A)** Top: Representative confocal image sequence showing the recruitement of EGFP-tagged wild-type OGG1 to 8-oxoG lesions generated by irradiation with a pulsed laser at 800 nm in the nucleus of OGG1 KO Hela cells. White arrowheads indicate the micro-irradiated line. Scale bar: 5 μm. Bottom: Recruitment kinetics of WT OGG1-EGFP at laser-induced damage measured from image sequences as shown at the top. Mean±SEM of 12 cells. **(B)** Normalized FCS autocorrelation curves measured of EGFP-tagged OGG1-WT and an EGFP-dimer (EGFP2) expressed in OGG1 KO Hela cells in the absence of DNA damage. Mean±SEM of 15 cells per condition. **(C)** Residence times of the EGFP-tagged constructs within the focal volume estimated from the fit of the autocorrelation curves shown in B. 15 cells per condition. **(D)** Sketch illustrating how base unstacking is detected by unquenching of the fluorescent nucleobase analog 2-aPu. **(E)** Stopped-flow 2-aPu fluorescence traces upon mixing the DNA substrate with OGG1-WT or with the substrate alone.

Altogether, our data draw a picture in which OGG1 rapidly transits on the DNA while searching for oxidized lesions, this transient binding leading to a local distortion of the DNA probably crucial for base inspection. In the following, we will use our analysis pipeline combining *in vitro* and live cell assays to interrogate the implication of conserved OGG1 residues in this rapid DNA sampling process.

### The NNN motif regulates the dynamics of OGG1 DNA sampling

Our recent report hinted at a key role of the highly conserved NNN motif (N149, N150 and N151) in the regulation of the DNA sampling process^21^. Yet, structural data indicates that these three Asn residues are differentially located relative to DNA during the lesion-seeking phase^17,24^, with N149 contacting simultaneously the guanine and the opposite cytosine, N151 interacting with the non-target strand, and N150 that does not display specific contacts with DNA (Figure 2A). To better understand the functions of this NNN motif, we studied more systematically the impact of mutating each specific Asn residue on OGG1 behavior in living cells.

**Figure 2.**
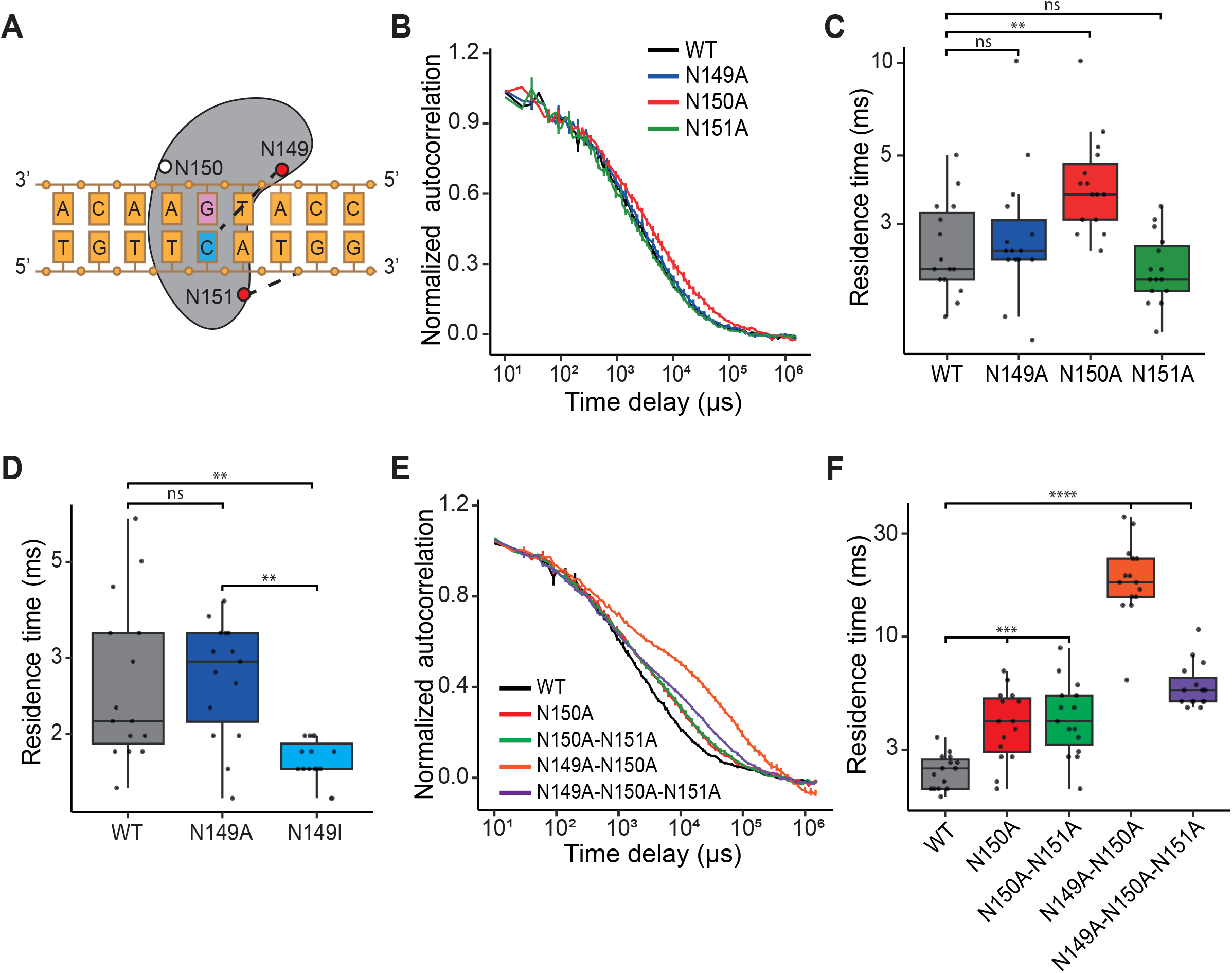
The NNN motif regulates OGG1 DNA sampling dynamics. **(A)** Sketch illustrating the interactions of the NNN motif with DNA during the lesion research phase according to published structural data. Residues in red establish DNA interactions shown with dashes. Residues that do not interact with DNA are shown in white. **(B–F)** Normalized FCS autocorrelation curves (B, E) and corresponding residence times within the confocal volume (C, D, F) for different EGFP-tagged OGG1 constructs in undamaged nuclei. Residence times were extracted from the fit of the FCS aucorrelation curves, 15 cells per condition.

First, we monitored the effect of individual mutations along this motif on the dynamics of DNA sampling in the absence of damage. Our FCS data revealed slower sampling dynamics for the N150A mutant while N149A and N151A were indistinguishable from WT, showing that the loss of N150 tend to specifically strengthen OGG1 association with DNA (Figure 2B,C). As N149 establishes hydrogen bonds with both the G:C and 8-oxoG:C pairs^17^, we also mutated this Asn into Ile to impair hydrogen bonding while maintaining steric hindrance. This N149I mutant showed accelerated dynamics compared to WT and N149A, suggesting reduced interaction with DNA (Figure 2D). To investigate potential compensatory mechanisms between neighboring residues, we also studied the impact of different combinations of mutations within the NNN motif. While combining N151A with N150A had no additional impact on OGG1 dynamics, the double mutant N149A/N150A (2NA) was dramatically slowed-down, a phenotype that was partially lost in the triple mutant N149A/N150A/N151A (3NA) (Figure 2E,F). Therefore, the NNN motif, and in particular residues N149 and N150, control OGG1 association with DNA during the lesion-seeking process.

### A coordinated role of the NNN motif and the wedge residue Y203 in DNA sampling and lesion detection

Because of the recent structural data reporting interactions between the wedge residue Y203 and the NNN motif^17^, we analyzed the impact of combining mutations on these residues on OGG1 ability to sample DNA during the lesion-seeking step. In undamaged nuclei, we found that, while the single Y203A mutation had no impact on OGG1 dynamics in living cells, it fully suppressed the trapping on the DNA observed for OGG1-N150A and OGG1-2NA (Figures S1A,B and 3A,B). These observations in living cells are in agreement with the analysis of the interactions between purified OGG1 and DNA oligonucleotides using electrophoretic mobility shift assay (EMSA), which showed that the increased affinity for DNA observed for the N150A and 2NA constructs compared to WT was lost upon addition of the Y203A mutation (Figure S1C-G). Conversely, the impaired interaction of OGG1-N149I with DNA reported in the cell nuclei could not be restored in constructs also bearing the Y203A mutation (Figure 3A,B).

**Figure 3.**
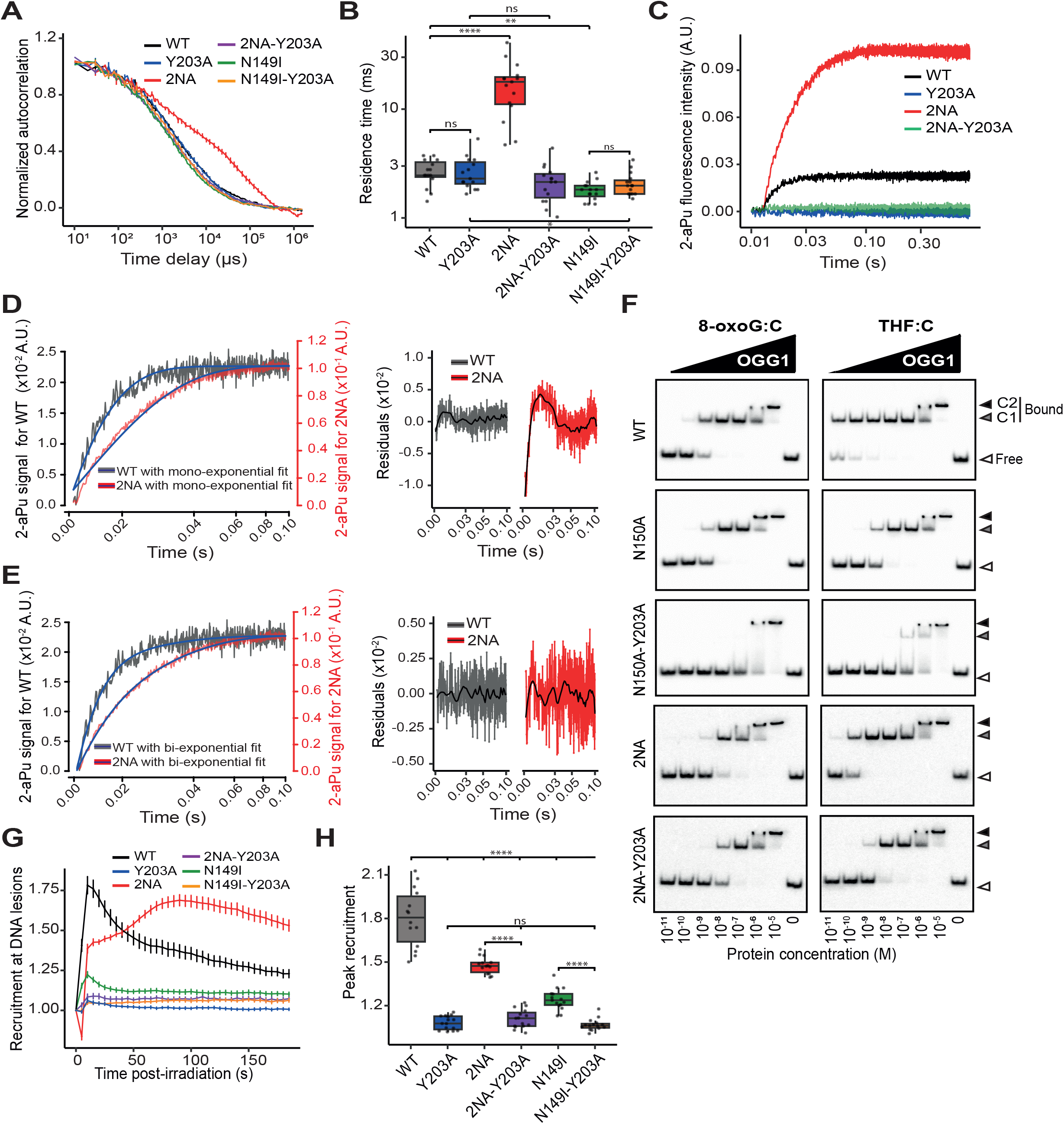
A coordinated role of the NNN motif and the wedge residue Y203 in DNA sampling and lesion detection. **(A-B)** Normalized FCS autocorrelation curves (A) and associated residence times (B) measured for EGFP-tagged OGG1 WT, Y203A, 2NA, 2NA-Y203A, N149I and N149I-Y203A in the absence of DNA damage. 15 cells per condition. **(C)** Stopped-flow 2-aPu fluorescence traces upon mixing the DNA substrate with OGG1-WT, Y203A, 2NA or 2NA-Y203A. **(D-E)** Stopped-flow 2-aPu fluorescence traces obtained with OGG1-WT and 2NA were fitted with a mono- (D) or a bi- (E) exponential model (black lines). Fit residuals are shown on the right. **(F)** Representative gel-shift blots showing the binding of purified OGG1 WT, N150A, N150A-Y203A, 2NA and 2NA-Y203A to radiolabeled DNA duplexes containing either an 8-oxoG:C or a THF:C pair. Complex C1 corresponds to the specific lesion recognition complex, with one protein bound per 8-oxoG:C probe. In contrast, complex C2 likely represents a non-specific interaction, where two OGG1 molecules bind to a single DNA duplex, independently of the presence of the lesion. **(G)** Recruitment kinetics of different OGG1 constructs at sites of DNA lesion measured from the image sequences as shown on figure S1H. **(H)** Recruitment peaks of the different OGG1 constructs extracted from curves shown in G. 14 to 15 cells per condition.

Next, we used fluorescence stopped flow to monitor double-helix distortion associated with DNA inspection by OGG1. In agreement with previous observations^14^, Y203A mutation completely suppressed 2-aPu unstacking, highlighting the central role of this residue in this process (Figure 3C). In contrast, mutations of Asn 149 and 150 increased 2-Pu fluorescence rise compared to WT. This could be due to the high affinity of OGG1-2NA for DNA but it may also reveal a more pronounced unstacking of 2-aPu. We also found that, while the rise of the 2-aPu fluorescence signal upon mixing with OGG1-wt could be fitted with a single exponential, the OGG1-2NA data rather followed a two-component exponential model, supporting a direct impact of Asn 149 and 150 on the kinetics of the base unstacking process (Figure 3D,E). Importantly, mutating Tyr 203 on the 2NA construct completely suppressed 2-Pu fluorescence rise similar to its effect on OGG1-WT, confirming that the impact of Asn 149 and 150 on double-helix distortion relies on Y203 wedge function (Figure 3C). Together with our data on the dynamics of OGG1 interactions with DNA, these results suggest that the NNN motif is an important regulator of OGG1 DNA sampling by controlling Y203-dependent base unstacking.

Finally, we also wondered whether the joint action of the NNN motif together with the Y203 wedge during base inspection impacted OGG1 ability to detect DNA lesions. EMSA revealed a reduced binding of N150A and 2NA mutants to both 8-oxoG and the AP-site analogue THF, which was impaired even further upon mutation of Y203 (Figure 3F). In cells, mutating the NNN motif had different impacts on OGG1 accumulation to DNA lesions. While N150A was indistinguishable from OGG1-WT (Figure S1I), the accumulation of the N149I mutant was drastically reduced and the 2NA construct showed delayed recruitment kinetics (Figures 3G,H and S1H). Furthermore, mutating the Y203 residue, alone or in association with other mutations in the NNN motif, completely abrogated OGG1 recruitment to the lesions (Figures 3G,H and S1H). Therefore, while residue Y203 is essential for lesion detection, the NNN motif is rather involved in the fine tuning of this process, thus affecting the amplitude and kinetics of OGG1 accumulation at sites of damage.

### Synergistic impacts of R154 and R204 on DNA sampling and recruitment to DNA lesions

OGG1 needs to specifically excise 8-oxoG paired with a cytosine to prevent G:C to T:A transversions. This discrimination step is thought to involve the two conserved residues Arg154 and Arg204. While both residues contacts and stabilize cytosine when 8-oxoG is extruded^24^, they rather interact with the non-target strand without direct contact with cytosine during the search phase^17^ (Figure 4A). To better understand the involvement of Arg 154 and 204 in the lesion-seeking process, we first studied the dynamics of the R154A and R204A single mutants as well as the R154A/R204A double mutant (2RA) *in vitro* and in living cells in the absence of inflicted damage. Using FCS, we found that the two single mutants OGG1-R154A and R204A both display accelerated dynamics compared to WT, this phenotype being even stronger with the 2RA mutant (Figure 4B,C). These results, in line with our EMSA data (FigureS1F,G), imply that Arg 154 and 204 both regulate OGG1 interactions with undamaged DNA. In contrast to the enhanced affinity of OGG1-2NA for DNA that relied on Y203 residue (Figure 3A), adding the Y203A mutation did not affect the dynamic of OGG1-2RA (Figure S2), suggesting that Arg 154 and 204 regulate OGG1 DNA sampling independently of Tyr 203. Using stopped-flow, we also found that OGG1-2RA was unable to promote double-helix distortion leading to 2-aPu unstacking (Figure 4D). Finally, EMSA revealed that the 2RA mutant lost its ability to bind to both 8-oxoG and THF (Figure 4E), and a cumulative impairment of OGG1 accumulation to sites of laser irradiation was observed in cells when mutating Arg 154 and 204 (Figure 4F-H). These data demonstrate that, in addition to their role in the detection of the estranged cytosine facing 8-oxoG, Arg 154 and 204 also act together at an early stage of the lesion-seeking process by controlling efficient DNA sampling and base unstacking.

**Figure 4.**
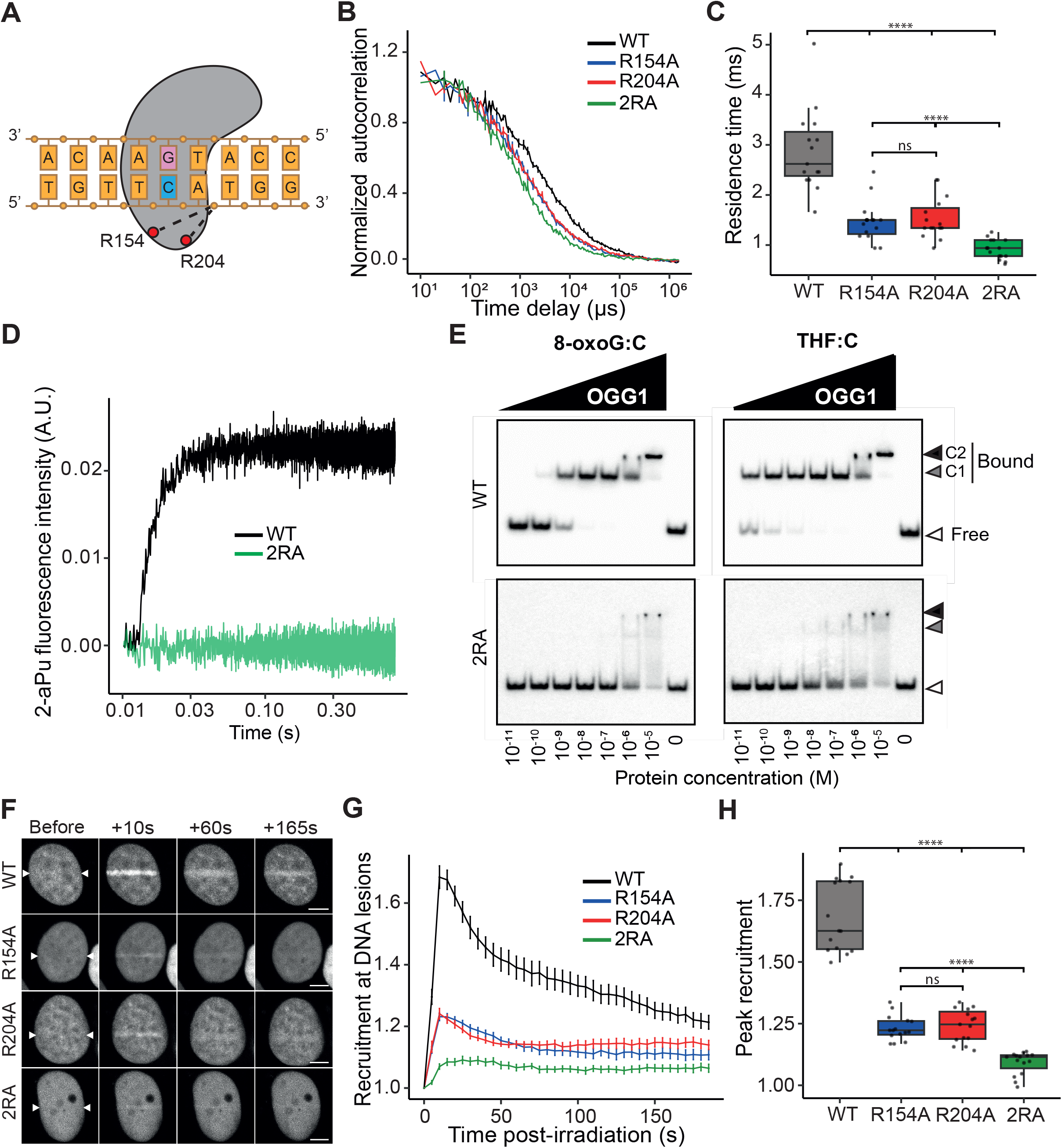
Synergistic impacts of R154 and R204 on DNA sampling and recruitment to DNA lesions. **(A)** Sketch illustrating the interaction of the two arginines 154 and 204 with DNA during the lesion-seeking phase according to published structural data. Residues in red establish DNA interactions shown with dashes. **(B-C)** Normalized FCS autocorrelation curves (B) and associated residence times (C) obtained for OGG1-WT, R154A, R204A and R154A/R204A (2RA) in the absence of DNA damage. 15 cells per condition. **(D)** Stopped-flow 2-aPu fluorescence traces upon mixing the DNA substrate with OGG1-WT and OGG1-2RA. **(E)** Representative gel-shift blots showing the binding of purified OGG1-WT, OGG1-2RA to radiolabeled DNA duplexes containing iether an 8-oxoG:C or a THF:C pair. The protein/DNA complexes C1 and C2 are defined in the caption of figure 3. The gel shown here for OGG1-WT is a duplication of the one already shown in figure 3F. **(F)** Representative image sequences of the accumulation of GFP-tagged OGG1 WT, R154A, R204A and 2RA at sites of laser irradiation in the nucleus of HeLa OGG1 KO cells. White arrowheads indicate the irradiated line. Scale bar: 5 μm. **(G-H)** Recruitment kinetics (G) and peak accumulation (H) of the different OGG1 constructs at DNA lesions derived from image sequences as shown in F. 15 cells per condition.

### Impairing OGG1 ability to efficiently sample DNA affects 8-oxoG excision activity

The ultimate goal of OGG1 is to efficiently clear 8-oxoG from the genome and we wondered how the contributions of the different residues to early stages of the base probing process investigated so far translates in terms 8-oxoG cleavage activity. We first monitored cleavage of 8-oxoG within oligos and found that mutants N150A and 2NA maintained a partial activity (Figure 5 A,B). Mutating Tyr 203, alone or in conjunction with mutations in the NNN motif, fully suppressed 8-oxoG cleavage, similar to what we observed for OGG1-2RA. To assess potential differential regulations within the intracellular context, we also monitored OGG1 activity in cells by quantifying 8-oxoG levels after oxidative DNA damage by potassium bromate (KBrO_3_). Immunofluorescence staining showed that, in contrast to OGG1-WT, OGG1-2RA was unable to restore basal levels of intranuclear 8-oxoG 4h after KBrO_3_ treatment (Figure 5C), thus confirming our *in vitro* results showing the loss of cleavage activity of this mutant. Altogether, these results show that mutating residues controlling lesion seeking ultimately impairs OGG1 ability to efficiently cleave its cognate lesion.

**Figure 5:**
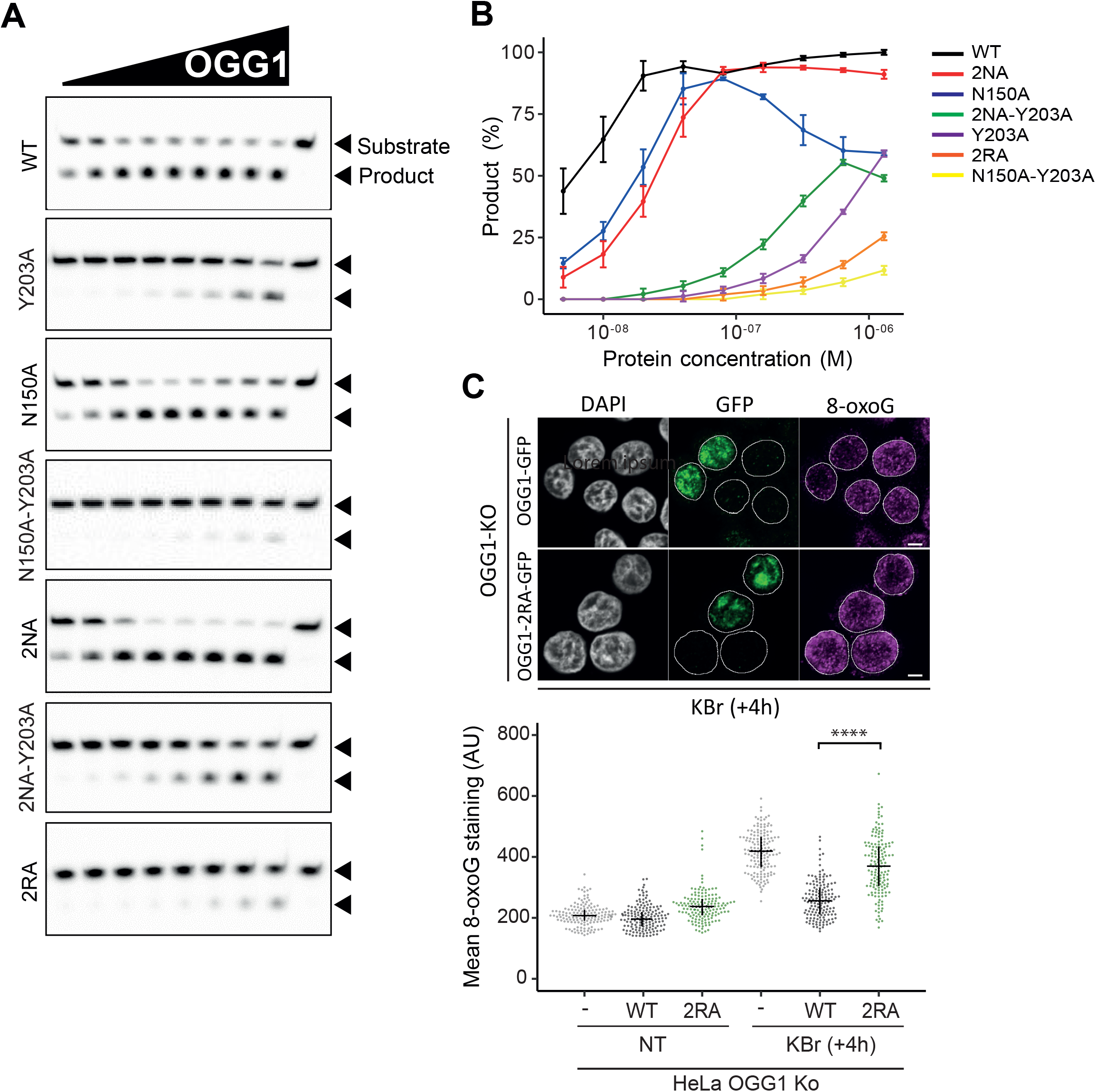
Impairing OGG1 ability to efficiently sample DNA affects 8-oxoG cleavage activity *in vitro* and in living cells. **(A)** Representative gels showing the cleavage of a radiolabeled oligonucleotide containing an 8-oxoG:C pair by OGG1-WT, 2NA and 2RA. **(B)** Relative amounts of cleaved product quantified from the gels shown in A. Each point represents mean±SD of three independent experiments. **(C)** Left: Representative images of the nuclei of OGG1 KO Hela cells expressing EGFP-tagged OGG1-WT or OGG1-2RA (green) fixed 4h after 45 min KBrO_3_ treatment, immunostained for 8-oxoG and colabeled with DAPI (gray). Scale bar: 5 µm. Right: 8-oxoG staining quantified from images as shown on the left before (NT) and 4 hours after 45 min KBrO_3_ treatment in OGG1 KO cells, untransfected or expressing EGFP tagged OGG1-WT or OGG1-2RA. 150 cells per condition.

## DISCUSSION

Similar to all factors searching for rare targets within the nucleus, OGG1 faces a major paradox: it must rapidly scan large fractions of the genome while maintaining high specificity for its cognate lesion. Pioneering theoretical work has shown more than 30 years ago that addressing this so-called speed-stability paradox requires that a given searching factor displays various modes of binding to the DNA, including some allowing diffusion along the double helix^25–27^. In line with these theoretical predictions, structural studies reported different conformations of the OGG1:DNA complex^17,24,28^. Nevertheless, beyond these structural snapshots, getting specific insights regarding transition kinetics between the binding states is thought to be crucial to understand how OGG1 samples the DNA, as shown for other nuclear factors^29–31^. Here, we combined *in vitro* and live cell assays to monitor the dynamic interactions of OGG1 and DNA and identify the key residues regulating this process.

Our work confirms the central contribution of Tyr 203 in base unstacking irrespectively of the presence of 8-oxoG^14^, an essential step that defines OGG1 ability to detect the lesions on naked DNA as well as in living cells. Yet, this residue does not regulate the dynamic association of OGG1 with DNA within the cell nucleus, in line with previous *in vitro* results showing that wedge residues do not impact the duration of glycosylase transits along undamaged DNA^12^. This implies that Tyr 203 only gently “pokes” the DNA to rapidly discriminates between G and 8-oxoG without requiring a lengthier complete extrusion. Importantly, our results reveal that the strength of this Y203-dependent poking, and consequently the speed of this initial base inspection step, is regulated by the NNN motif. These conserved residues establish a network of contacts with both Y203 and the DNA backbone, irrespectively of the oxidation status of the guanine in OGG1:DNA complex conformations thought to initiate the lesion detection process^17,24^. Together, our results and these structural insights suggest that the NNN motif might impact OGG1 binding to DNA and base unstacking in two alternative ways (Figure 6). First, it may act as a “return spring” to control the insertion of the Y203 poking tip within the double helix. Alternatively, the Asn stretch might regulate local DNA bending, which in turns would modulate Y203-dependent base unstacking. These two regulatory mechanisms would favor a transient semi-extruded base conformation displaying limited distortion of the double helix to avoid a local collapse of the DNA structure leading to OGG1 trapping. While not strictly needed for 8-oxoG detection and cleavage, this NNN spring would then be crucial for rapid DNA probing, a feature that appears particularly critical for the detection of rare 8-oxoG lesions which frequency is as low as few oxidized bases per 10^6^ guanines in unchallenged conditions^32^.

**Figure 6:**
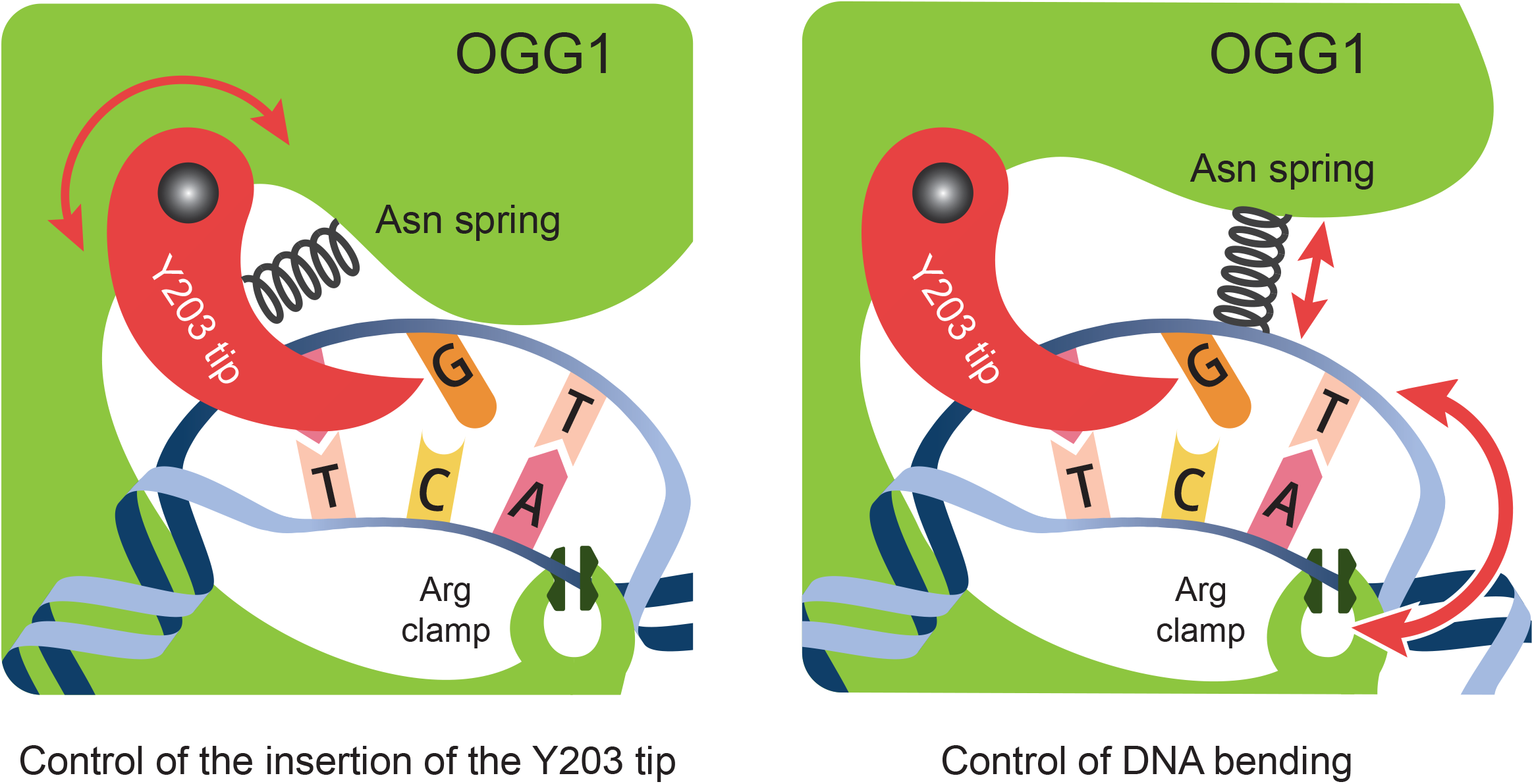
Possible mechanisms by which the NNN motif and Arg 154 and 204 act in coordination with Y203 during lesion-seeking by OGG1. **Left :** The NNN motif can act as a “return spring” to control the insertion of the Y203 tip in the double helix. **Right :** The NNN motif could regulate local DNA bending, which in turns would modulate Y203-dependent base unstacking. In these two models, Arg 154 and 204 act as a clamp on the non-target strand to regulate transient association with DNA.

Besides the coordinated action of Y203 and the NNN motif during G and 8-oxoG unstacking and extrusion, our results reveal that arginines 154 and 204 are also crucial for these early stages of OGG1 base probing. While these two residues are well known to contribute to the recognition of the estranged cytosine^24,33^ upon 8-oxoG extrusion, they also establish direct contacts with the phosphate backbone of the non-target strand before the extrusion stage^17^. In line with these structural data, our findings show that Arg 154 and 204 both contribute to OGG1 transient binding to DNA irrespectively of the presence of 8-oxoG and, by clamping the non-target strand (Figure 6), are essential for Y203-dependent base unstacking. Therefore, these two conserved residues are not only important to ensure OGG1 specificity for 8-oxoG:C pairs, but also to initiate the base probing process.

Altogether, our data reveal that OGG1 base probing is initiated by a tight coordination between several conserved amino-acids encircling the double-helix. These residues finely tune OGG1 binding to DNA to ensure that this association is stable enough for base interrogation but also to prevent an excessive engagement that would be detrimental to fast scanning dynamics. Together with the wedge residue Y203, the NNN motif as well as Arg 154 and 204 also control intrahelical base destabilization irrespectively of the oxidized form of guanine, a prerequisite to the subsequent discrimination between G and 8-oxoG. This cooperative function of multiple residues appears central to enable rapid exhaustive DNA sampling within the nucleus while maintaining high specificity for 8-oxoG:C pairs. Besides 8-oxoG clearing, this fine regulation of DNA binding kinetics might also be crucial for OGG1 contribution to the inflammatory response. In line with this hypothesis, it was recently proposed that the proinflammatory impact of the frequent OGG1 variant S326C was accounted by its increased binding at 8-oxoG containing inflammatory gene promoters^34^. Future work will be needed to assess the contributions of the NNN motifs as well as Arg 154 and 204 in these processes.

## MATERIAL AND METHODS

### Plasmids

All plasmids used in this study are listed in Table S1. The following plasmids were previously described: OGG1-L1-EGFP, OGG1(N149A-N150A)-L1-EGFP, OGG1(Y203A)-L1-EGFP, His-OGG1(Y203A) and His-OGG1(N149A-N150A)^21^, EGFP2^35^. All OGG1 plasmids refer to the α isoform of human OGG1. Point mutations in OGG1-L1-EGFP were generated using site-directed mutagenesis with primers listed in Table S2. The plasmids OGG1(N149I-Y203A)-L1-EGFP, OGG1(N150A-Y203A)-L1-EGFP and OGG1(R154A-R204A)-L1-EGFP were generate in 2 steps. All constructs were validated by DNA sequencing. Rescriction enzymes and DNA polymerases were from Thermo, antibiotics from euromedex and primers from Sigma. The mutant constructions of His-tagged OGG1 were generated as previously described^21^. Site-directed mutagenesis was performed on the recombinant pET30a-hOGG1 plasmid using the Quick-Change kit (Agilent) to introduce point mutations, with primers listed in Table S2. All constructs were validated by DNA sequencing. Restriction enzymes and DNA polymerases were from New England Biolabs, antibiotics from Sigma-Aldrich and primers from Eurogentec.

### Cell culture

Routine cell culture was performed in DMEM (Sigma) supplemented with 10% fetal bovine serum (FBS), 100 µg/mL penicillin, 100 U/mL streptomycin and maintained at 37 °C in a 5% CO_2_ incubator. HeLa cells knocked-out for OGG1 were generated previously^21^. For all live-cell experiments, cells were cultured in 8-well Imaging Chamber CG (Zell-Kontakt) and transfected with the specified plasmid 24 to 48 hours before imaging, using the X-TremeGENE HP (Roche) transfection reagent in accordance with the manufacturer’s instructions. Just before imaging, growth medium was replaced with CO_2_-independent imaging medium (Phenol Red-free Leibovitz’s L-15 medium (Life Technologies) supplemented with 20% fetal bovine serum, 2 mM glutamine, 100 µg/mL penicillin and 100 U/mL streptomycin). All experiments presented in this work were performed on unsynchronized cells.

### Protein purification

For stopped-flow measurements, the plasmids pET30a-OGG1, pET30a-OGG1-Y203A, pET30a-OGG1-N149A-N150A, pET30a-OGG1-N149A-N150A-Y203A, and pET30a-OGG1-R154-R204 encoding the 6×His-tagged proteins OGG1-WT, OGG1-Y203A, OGG1-N149A-N150A, OGG1-N149A-N150A-Y203A, and OGG1-R154-R204 were transformed into *E. coli* BL21 (DE3). Cultures were grown at 37 °C with 1 mM IPTG added when optical density reached 0.6, followed by an additional 4-hour incubation at 37 °C. Cells were harvested by centrifugation and resuspended in lysis buffer (20 mM HEPES, pH 7.6, 500 mM NaCl, 5% glycerol, 20 mM imidazole pH 8, 1 mM DTT) followed by sonication. The lysate was clarified by centrifugation at 20,000 × g for 1 hour at 4 °C. The supernatant was loaded onto a 5 mL HisTrap™ HP column (Cytiva), precharged with nickel, according to the manufacturer’s instructions, and equilibrated with the same lysis buffer. The column was washed with washing buffer (20 mM HEPES, pH 7.6, 2 M NaCl, 5% glycerol, 1 mM DTT), and proteins were eluted using a linear gradient of elution buffer (20 mM HEPES, pH 7.6, 150 mM NaCl, 5% glycerol, 300 mM imidazole pH 8, 1 mM DTT), up to 60% of the elution buffer. Fractions containing purified OGG1-WT and the corresponding mutant constructs were pooled, concentrated, and stored in buffer containing 350 mM NaCl, 20 mM HEPES pH 7.5, 1 mM DTT, 5% glycerol at -80 °C. For electrophoretic mobility shift assay (EMSA) and cleavage activity assays, His-tagged OGG1-WT and the mutant variants N150A, N149A-N150A, Y203A, N150A-Y203A, N149A-N150A-Y203A, and R154A-R204A were produced and purified as previously described^21^. Briefly, after expression in bacteria, the 6×His tag was removed by enterokinase digestion. The digestion mixture was then applied to a benzamidine column (GE Healthcare) followed by a HisTrap™ column (GE Healthcare). The protein was further purified by cation exchange chromatography using a POROS™ HS20 column. Elution fractions containing the protein of interest were pooled and subjected to size-exclusion chromatography on a Superdex 75 column (GE Healthcare). The purified proteins were concentrated and stored at –80 °C.

### Stopped-flow fluorescence measurements

Stopped-flow data were recorded on a BioLogic SFM-300, MOS-250 instrument. The dead time of the stopped-flow device was 0.125 ms. All experiments were carried out at 25°C in a buffer containing 50 mM Tris–HCl, pH 7.5, 75 mM KCl, 1 mM EDTA, 1 mM DDT and 9% glycerol. The temporal evolution of 2-aPu fluorescence were monitored at excitation and emission wavelengths of 312 and 370 nm, respectively. OGG1 contained in one syringe was rapidly mixed with the double-stranded oligo bearing 2-aPu contained in the second syringe. The concentration of OGG1 after mixing in all experiments was 2.0 μM, whereas that of the substrate was 1.0 μM. The curves are from a characteristic experiment among at least three independent repetitions and each curve is the average of 10-20 shots. We used the R software (https://www.r-project.org/) to fit the fluorescence traces using the following exponential model:

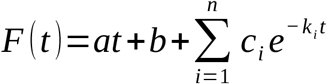

where *c*_*i*_ and *k*_*i*_ correspond to the amplitude and rate constant of the exponential component, respectively, a is the amplitude of the linear component and b is the offset. *n* was equal to 1 or 2 for the mono- or bi-exponential models, respectively.

### Electrophoretic mobility shift assay (EMSA)

EMSA was used to assess the ability of OGG1-WT and the mutant variants OGG1-N150A, OGG1-N150A-Y203A, OGG1-N149A-N150A-Y203A, and OGG1-R154A-R204A to bind the oligonucleotides listed in Table S2. 0.1 nM of 5′-[^32^P]-labeled 24-mer DNA duplexes were incubated at 4 °C for 30 min, either alone or with the indicated protein concentrations, in binding buffer. Samples were then resolved on 10% native polyacrylamide gels. After drying, the gels were exposed using a Typhoon molecular imager (Amersham) for autoradiography and quantified with ImageQuant software. Triplicate EMSA titrations were performed to determine the apparent dissociation constants (K_Dapp_). Finally, using Origin software, binding curves representing the relative quantities of DNA/protein complexes (C) as a function of protein concentration (P) were adjusted with a non-linear regression following Hill’s equation:

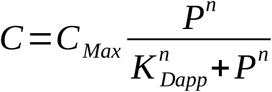

where *C*_*Max*_ denotes the maximal level of DNA/protein complex, *K*_*Dapp*_ is the apparent dissociation constant, and *n* corresponds to the Hill coefficient.

### OGG1 DNA glycosylase activity

Glycosylase activity assays for OGG1-WT and its various mutants were performed as previously described^21^. Briefly, 20 nM of 24-mer [8-oxoG:C] DNA duplexes labeled at the 5′ end with [^32^P] were incubated at 37 °C for 15 minutes, either alone or with varying concentrations of OGG1 (WT or mutant variants). Reactions were terminated by the addition of 0.2 M NaOH followed by a 5-minute incubation at 50 °C, and then incubated for 3 minutes at 75 °C in a formamide loading dye buffer. Reaction products were separated by 20% denaturing polyacrylamide gel electrophoresis (PAGE), visualized by autoradiography using a Typhoon scanner (Amersham), and quantified with ImageQuant software.

### Laser micro-irradiation

The recruitment dynamic of OGG1 and its mutant variants was analyzed using a Zeiss LSM 880 confocal microscope equipped with a 40×/1.2 NA C-Apochromat water-immersion objective. EGFP was excited at 488 nm, and emission was detected in the 500–550 nm range via a GaAsP detector. The laser power has been adjusted to limit photobleaching, and images have been acquired at a pixel resolution of 80 nm. Localized DNA damage (8-oxoG lesions) was induced by targeting a defined nuclear region (12 µm × 0.64 µm) using an 800 nm Ti:Sapphire femtosecond laser (Mai Tai HP, Spectra Physics). Time-lapse imaging was performed every 5 s over a 200 s period to monitor protein accumulation kinetics at sites of damage. To void bias due to protein expression variability, cells with similar nuclear EGFP signals were analyzed across all conditions. All experiments were conducted at 37 °C using a heating chamber.

### Image analysis

To quantify protein recruitment kinetics at sites of damage, the mean intensities within the damaged region (*I*_*d*_), the nucleus (*I*_*n*_), and the background outside of the cell (*I*_*bg*_) were measured by segmentation using ImageJ. Protein accumulation at sites of damage (*PA*_*d*_) was then calculated as:

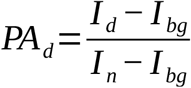

The peak recruitment was considered to be the maximum intensity value within the first 30 seconds after damage induction.

### Fluorescence correlation spectroscopy data

FCS measurements were carried out on a Zeiss LSM 880 confocal microscope using a 40×/1.2 NA C-Apochromat water-immersion objective. EGFP fluorescence was excited at 488 nm, and emitted photons were collected in the 500–550 nm range and recorded in photon-counting mode using a GaAsP detector. Laser intensity was adjusted to minimize photobleaching. Each FCS acquisition lasted 20 s to reduce the noise in the autocorrelation curves. Raw photon traces were processed with the Fluctuation Analyzer 4G software, applying a detrending frequency of 0.250 Hz to correct for slow signal fluctuations^36^. Autocorrelation curves were then fitted using a standard diffusion model:

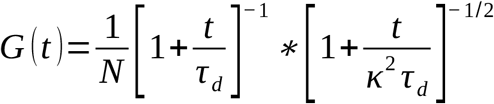

where *N* is the number of fluorescent protein within the confocal volume, *τ*_*D*_ is the characteristic residence time within the confocal volume and *k* is the structural parameter, which is fixed at 6.

### Cell nucleus 8-oxoG clearance assay

For visualization of 8-oxoG nuclear levels, HeLa OGG1 KO cells expressing OGG1(WT)-GFP or OGG1(2RA)-GFP were grown on µ-Slide 8-well glass bottom dishes (Ibidi, 80827). Oxidative DNA damage was induced by adding 40 mM potassium bromate (KBrO3; Sigma) diluted in DPBS (Sigma) for 45 min at 37°C, followed by 4h recovery in culture medium. Cells were fixed in acetone:methanol (1:1) and air dried. Cells were hydrated for 15 min in PBS, and DNA was denatured by incubating cells in 2N HCl for 45 min at room temperature. Cells were washed three times in PBS, neutralized with 50 mM Tris–HCl pH 8.8 for 5 min and permeabilized at room temperature in PBS–0.1% Triton for 10 min. Cells were incubated in blocking solution (PBS, 0.1% Triton, 3% BSA, 1% normal goat serum) at 37°C for 1 h and subsequently incubated for 1 h at 37°C with antibodies against 8-oxoG (ab48508, abcam) and GFP (632592, Clontech) diluted in blocking solution at 1:2000 and 1:1000, respectively. Cells were then washed three times for 10 min in PBS–0.1% Triton and incubated with an anti-Mouse antibody coupled to Alexa-594 (A11005, Thermofisher) and an anti-Rabbit antibody coupled to Alexa-488 (A11008, Thermofisher), both diluted at 1:1000 in blocking solution for 1 h at 37°C. Nuclear DNA was counterstained with 1 μg/ml DAPI. Cells were imaged with a Nikon A1 confocal microscope, using a ×60 oil immersion objective with a numerical aperture of 1.3 and analysis was performed with ImageJ. Fluorescence quantifications were performed by segmenting nuclei based on the DAPI signal, and applying the resulting masks to the 8-oxoG and EGFP channels to extract mean fluorescence intensities per nucleus. For the OGG1-WT and OGG1-R154A-R204A conditions, only EGFP-positive cells were included in the quantification of 8-oxoG signals.

### Statistics

Data analysis and figure generation were conducted using R software. In the boxplots, the boxes represent the interquartile range (25th to 75th percentiles), with the horizontal line indicating the median. Whiskers extend up to 1.5 times the interquartile range. For recruitment and FCS curves, data are presented as mean ± SEM. All results shown are from a representative experiment out of at least three independent replicates. Statistical comparisons were performed using two-sided, unpaired Student’s t-tests with unequal variance assumptions. Significance levels are indicated as follows: *p < 0.05; **p < 0.01; ***p < 0.001; ****p < 0.0001; n.s. = not significant.

## ACKNOWLEDGMENTS

We thank Microscopy-Rennes Imaging Center (BIOSIT, Univ Rennes 1), a member of France-BioImaging supported by the French National Research Agency (ANR) (grant no ANR-10-INBS-04), for access to optical microscopy and Stéphanie Dutertre, Xavier Pinson and Gilles Le Marchand for their assistance on the microscopes. We thank Christophe Tascon for his technical support during protein purification as well as Emmanuel Giudice and J. Pablo Radicella for advices and critical reading of the manuscript. S.H. and A.C. received joint financial support from the Agence Nationale de la Recherche [PRC 2023, CE12-0025, OxiREPTRA]. S.H. and B.C. were also funded by a joint grant from the Ligue contre le Cancer du Grand-Ouest [committees 37 and 45]. O.D.A.’s PhD fellowship was funded by the Région Bretagne, the CEA and the Ligue Nationale Contre le Cancer.

