## Supplemental info for "A coordinated haptic mechanism ensures efficient DNA sampling by the 8-oxoguanine glycosylase OGG1"

**This PDF file includes:**

Supplementary Table S1 and S2

Supplementary Figures S1 to S2

### SUPPLEMENTARY TABLES

**Table S1. List of plasmids used in this study**

| Plasmid | Source |
|---|---|
| OGG1-L1-GFP | D'augustin *et al.*, 2023 |
| OGG1(N149A)-L1-GFP | This study |
| OGG1(N149I)-L1-GFP | This study |
| OGG1(N150A)-L1-GFP | This study |
| OGG1(N151A)-L1-GFP | This study |

| | |
|---|---|
| OGG1(N150A-N151A)-L1-GFP | This study |
| OGG1(N149A-N150A)-L1-GFP | D'augustin *et al.*, 2023 |
| OGG1(N149A-N150A-N151A)-L1-GFP | This study |
| OGG1(Y203A)-L1-GFP | D'augustin *et al.*, 2023 |
| OGG1(N150A-Y203A)-L1-GFP | This study |
| OGG1(N149I-Y203A)-L1-GFP | This study |
| OGG1(N149A-N150A-Y203A)-L1-GFP | This study |
| OGG1(R154A)-L1-GFP | This study |
| OGG1(R204A)-L1-GFP | This study |
| OGG1(R154A-R204A)-L1-GFP | This study |
| GFP2 | Gift from Jan Ellenberg (Euroscarf P30623), Bancaud et al., 2009 |
| His-OGG1 | Le Meur et al., 2015 |
| His-OGG1(Y203A) | D'augustin *et al.*, 2023 |
| His-OGG1(N150A) | This study |
| His-OGG1(N150A-Y203A) | This study |
| His-OGG1(N149A-N150A) | D'augustin *et al.*, 2023 |
| His-OGG1(N149A-N150A-Y203A) | This study |
| His-OGG1(R154A-R204A) | This study |

**Table S2. List of oligonucleotides used in this study**

| Oligonucleotide | Sequence (5' -> 3') |
|---|---|
| OGG1_N149A_Fwd | CTGTTCCTCCGCCAACAACATC |
| OGG1_N149A_Rev | GATGTTGTTGGCGGAGGAACAG |
| OGG1_N149I_fwd | CTGTTCCTCCATCAACAACATC |
| OGG1_N149I_rev | GATGTTGTTGATGGAGGAACAG |
| OGG1_N151A_Fwd | CTCCAACAACGCCATCGCCC |
| OGG1_N151A_Rev | GGGCGATGGCGTTGTTGGAG |
| OGG1_Y203A_Rev | GTAACGGGCACGAGCGCCCAGGCCCAGC |
| OGG1_Y203A_Fwd | GCTGGGCCTGGGCGCTCGTGCCCGTTAC |
| OGG1_R154A_Fwd | CAACAACATCGCCGCCATCACTGGCATGG |
| OGG1_R154A_Rev | CCATGCCAGTGATGGCGGCGATGTTGTTG |
| OGG1_R204A_Fwd | GGCCTGGGCTATGCTGCCCGTTACG |

| | |
|---|---|
| OGG1_R204A_Rev | CGTAACGGGCAGCATAGCCCAGGCC |
| OGG1_N150A_Fwd | TCTCTTTTATCTGTTCCTCCAACGCCAACATCGCCCGCATCACTGGCATGGTG |
| OGG1_N150A_Rev | CACCATGCCAGTGATGCGGGCGATGTTGGCGTTGGAGGAACAGATAAAAGAGA |
| OGG1_N150A_N151A_Fwd | TCTCTTTTATCTGTTCCTCCAACGCCGCCATCGCCCGCATCACTGGCATGGTG |
| OGG1_N150A_N151A_Rev | CACCATGCCAGTGATGCGGGCGATGGCGGCGTTGGAGGAACAGATAAAAGAGA |
| OGG1_N150A_N151A_Rev | TCTCTTTTATCTGTTCCTCCGCGGCCGCCATCGCCCGCATCACTGGCATGGTG |
| OGG1_N149A-N150A_N151A_Fwd | TCTCTTTTATCTGTTCCTCCGCGGCCGCCATCGCCCGCATCACTGGCATGGTG |
| OGG1_N149A-N150A_N151A_Rev | ACCATGCCAGTGATGCGGGCGATGGCGGCCGCGGAGGAACAGATAAAAGAGA |
| Histag_OGG1_N150A_Fwd | CTTTTATCTGTTCCTCCAACGCGAACATCGCCCGCATCACTGG |
| Histag_OGG1_N150A_Rev | CCAGTGATGCGGGCGATGTTCGCGTTGGAGGAACAGATAAAAG |
| Histag_OGG1_Y203A_Fwd | GCCCGTGCCCGTTACGTG |
| Histag_OGG1_Y203A_Rev | CGGGCACGGGCGCCCAGGCCCAGC |
| Histag_OGG1_N149A-N150A_N151A_Fwd | GCGGCCAACATCGCCCGCATCAC |
| Histag_OGG1_N149A-N150A_N151A_Rev | CGATGTTGGCCGCGGAGGAACAGATAAAAGAGAAAAGGC |
| Histag_OGG1_R154A_Fwd | CCAACAACAACATCGCCGCGATCACTGGCATGGTGG |
| Histag_OGG1_R154A_Rev | CCACCATGCCAGTGATCGCGGCGATGTTGTTGTTGG |
| Histag_OGG1_R204A_Fwd | GCTGGGCCTGGGCTATGCGGCCCGTTACGTGAGTGC |
| Histag_OGG1_R204A_Rev | GCACTCACGTAACGGGCCGCATAGCCCAGGCCCAGC |
| 5’_[32P]_labeled_undamaged_or_8oxoG | CTGATCGATGAC**X**CCTGACATGAT |
| Complementary G(C)G for 5’-[32P]-labeled | ATCATGTCAGGCGTCATCGATCAG |
| 2-aPu_labeled_Fwd | CTCT[2-aPu]CTTCC |
| Complementary G(C)G for 2-aPu-labeled | GGAAGCGAGAG |

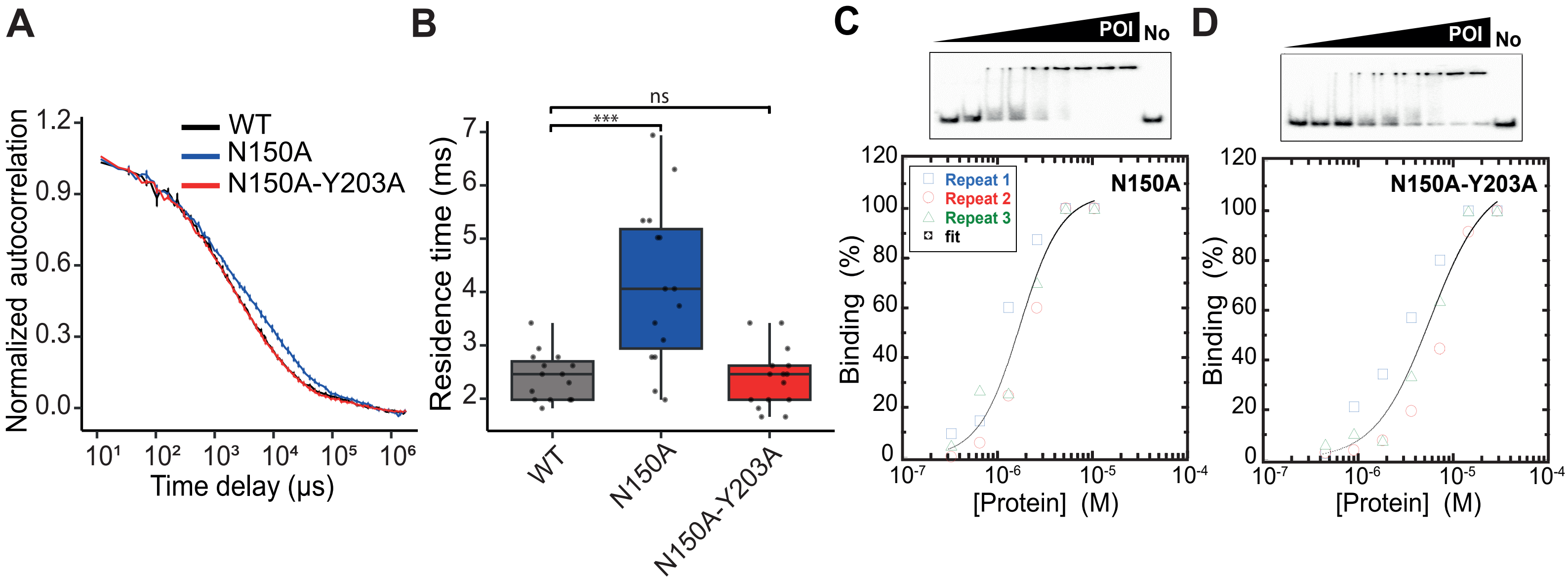


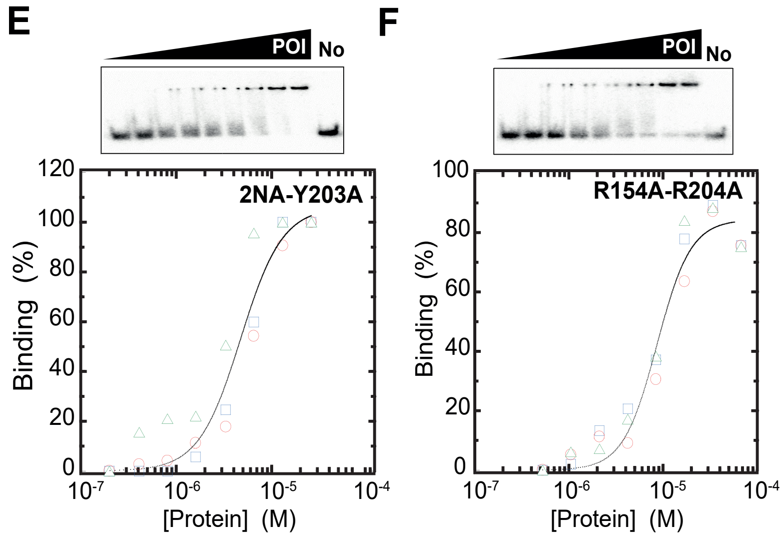


G

| Protein | $K_{Dapp}$ (mM) G:C |
|---|---|
| WT | 2.22 ± 0.22* |
| Y203A | 1.97 ± 0.27* |
| 2NA | 0.60 ± 0.04* |
| N150A | 1.73 ± 0.20 |
| N150A-Y203A | 5.74 ± 1.20 |
| 2NA-Y203A | 8.61 ± 0.73 |
| 2RA | 4.72 ± 0.65 |

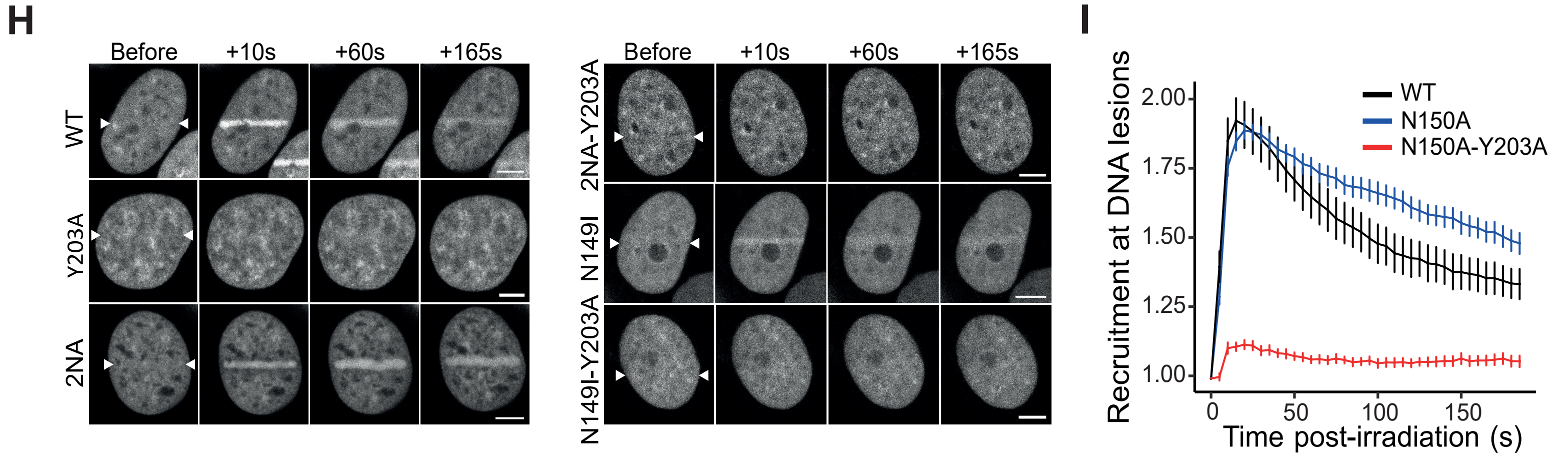


**Figure S1: The addition of the Y203A mutation suppresses the enhanced affinity of the N150A mutant for undamaged DNA and impairs its accumulation to DNA lesions. (A-B)** Normalized FCS autocorrelation curves (A) and associated residence times (B) measured for GFP-tagged OGG1-WT, N150A and N150A-Y203A in the absence of DNA damage. 15 cells per condition. **(C-F)** Binding of purified OGG1-N150A (C), OGG1-N150A-Y203A (D), OGG1-2NA-Y203A (E) and OGG1-R154A-R204A (F) to a 24-mer [G:C] duplex were analysed by EMSA. For each panel the upper part shows a representative gel autoradiography and the bottom part is a quantification of the relative amount of OGG1:DNA complex. Each point is the mean of three independent experiments. **(G)** Apparent dissociation constants estimated as the protein concentrations corresponding to 50% binding based on the titration curves shown on panels C to F. The dissociation constants for OGG1-WT, OGG1-Y203A and OGG1-2NA were estimated previously in D'Augustin et al., 2023. **(H)** Representative image sequences of the accumulation of EGFP-tagged OGG1-WT, Y203A, 2NA, 2NA-Y203A, N149I and N149I-Y203A at sites of laser irradiation in the nucleus of HeLa OGG1 KO cells. White arrowheads indicate the irradiated line. Scale bar: 5 µm. **(I)** Recruitment kinetics of OGG1-WT, N150A and N150A-Y203A at sites of laser irradiation. 12 cells per condition.

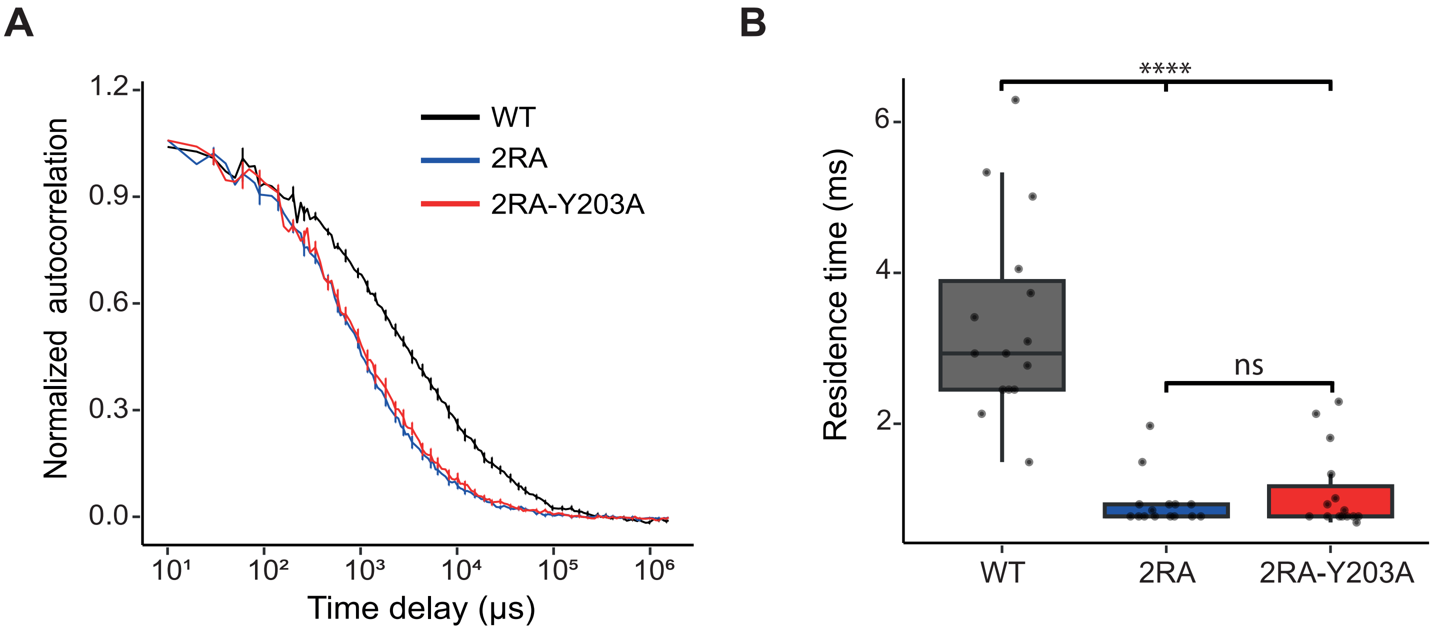


**Figure S2: The introduction of the Y203A mutation does not impact the reduced association of the 2RA mutant with undamaged DNA. (A-B)** Normalized FCS autocorrelation curves (A) and associated residence times (B) measured for OGG1-WT, 2RA and 2RA-Y203A in the absence of DNA damage. 15 cells per condition.
